# DNAscan: a fast, computationally and memory efficient bioinformatics pipeline for the analysis of DNA next-generation-sequencing data

**DOI:** 10.1101/267195

**Authors:** A Iacoangeli, A Al Khleifat, W Sproviero, A Shatunov, AR Jones, R Dobson, SJ Newhouse, A Al-Chalabi

**Affiliations:** Department of Biostatistics and Health Informatics, King’s College London, London, UK; Maurice Wohl Clinical Neuroscience Institute, King’s College London, London, UK; Farr Institute of Health Informatics Research, UCL Institute of Health Informatics, University College London, London, UK; National Institute for Health Research (NIHR) Biomedical Research Centre and Dementia Unit at South London and Maudsley NHS Foundation Trust and King’s College London London, UK.

**Keywords:** NGS, Miseq, WGS, WES, Bioinformatics, Variant calling, Structural variants, Repeat expansions, Virus and Bacteria identification

## Abstract

The generation of DNA Next Generation Sequencing (NGS) data is a commonly applied approach for studying the genetic basis of biological processes, including diseases, and underpins the aspirations of precision medicine. However, there are significant challenges when dealing with NGS data. A huge number of bioinformatics tools exist and it is therefore challenging to design an analysis pipeline; NGS analysis is computationally intensive, requiring expensive infrastructure which can be problematic given that many medical and research centres do not have adequate high performance computing facilities and the use of cloud computing facilities is not always possible due to privacy and ownership issues. We have therefore developed a fast and efficient bioinformatics pipeline that allows for the analysis of DNA sequencing data, while requiring little computational effort and memory usage. We achieved this by exploiting state-of-the-art bioinformatics tools. DNAscan can analyse raw, 40x whole genome NGS data in 8 hours, using as little as 8 threads and 16 Gbs of RAM, while guaranteeing a high performance. DNAscan can look for SNVs, small indels, SVs, repeat expansions and viral genetic material (or any other organism). Its results are annotated using a customisable variety of databases including ClinVar, Exac and dbSNP, and a local deployment of the gene.iobio platform is available for an on-the-fly result visualisation.

## Introduction

Next Generation Sequencing (NGS) technologies play a key role in human genetic research. The effort needed to sequence a whole genome has reduced from about 15 years of work at a cost of $3 billion in 2003 [1] to hours for ~$1000 in 2017 (https://www.veritasgenetics.com/mygenome).

Producing sequencing data, whether it is whole genome sequencing (WGS), whole exome sequencing (WES) or targeted gene panels, is common practice for the study of the genetic basis of biological processes. In biomedical research, NGS data are widely used to investigate the genetic causes of disease, allowing for the study of genomic variants including single nucleotide variations, small insertions or deletions of a few bases, as well as structural variants.

On a large scale, international collaborations are forming sequencing consortia to study the genetic landscape of thousands of individuals. Examples of such consortia are Project MinE [2] and The Cancer Genome Atlas (TCGA) [3]. Project MinE is an international consortium seeking to obtain sequencing data from 15,000 Amyotrophic Lateral Sclerosis cases and 7,500 matched controls. TCGA is a rich dataset of sequencing data of over 11,000 individuals affected by 33 different tumour types. On an individual scale, NGS data are also being investigated for their use in diagnostic medicine and so called Precision Medicine [4, 5], whose aim is to tailor medical treatments to patient genetics.

There are several practical challenges when processing NGS data. For example, 40x WGS data for one sample produced on the Illumina Hiseq 2000, one of the most popular sequencers, is about 400 gigabytes in its raw format (fastq format) [6]. This size can be reduced to approximately one fourth when the data is compressed down to about 100 gigabytes, using lossless formats such as fastq.gz (gzip-compressed version of fastq) and bam [7]. Such big files are not easy to handle for the average non-specialised scientist or lab, since they require sophisticated tools, bioinformatics skills and high performance computing for their analysis. Indeed, as an example, consider mapping one of these files, typically about 1 billion 150-base-pair long reads, to the human genome, a key process in the analysis of WGS data. Assuming that a standard midrange desktop computer with 4 CPUs and 16 Gb RAM is used with The Burrow Wheeler Aligner (BWA) [8], probably the most widely used mapper, aligning this data to the human reference genome would take about 1 day, and this would only be the first step of an NGS data analysis pipeline. Faster mappers exist. For example SNAP [9] would only take about 4 hours to complete the same job, using the same number of CPUs, but it requires about 65Gb RAM, making it an unsuitable choice if large memory High-Performance-Computing (HPC) facilities are not available.

For big collaborations and projects, collecting NGS data from thousands of individuals, powerful and expensive HPC facilities, as well as highly specialised staff are needed. To handle such data, the Project MinE consortium makes use of SURFsara, the Dutch national HPC facility, and the TCGA invested millions of dollars in HPC infrastructure and e-infrastructure (https://cancergenome.nih.gov) making use of, among others, Amazon Web Services (AWS) and Seven Bridges (https://www.sevenbridges.com).

Further challenges are represented by the large number of bioinformatics tools available for NGS analysis. Omictools [10], a web database where most available tools are listed and reviewed, lists over 7000 bioinformatics NGS tools, and given the great interest in this field, new tools are frequently released. Therefore, designing a bioinformatics pipeline for the analysis of NGS data, taking into account both the available computing facilities and the study aim, is not trivial, and requires specialised expertise.

Here we describe DNAscan, an extremely fast, accurate and computationally light bioinformatics pipeline for the analysis, annotation and visualisation of DNA NGS data. DNAscan is designed to provide a powerful and easy-to-use tool for applications in biomedical research and diagnostic medicine, at minimal computational cost. DNAscan can analyse 40× WGS data in 8 hours using 8 threads and 16 Gb RAM and WES data in 1 hour using 4 threads and 10 Gb of RAM, enabling the processing of NGS data to be carried out on most midrange computers and the minimisation of computational costs. The pipeline can detect SNVs, small indels, SVs, repeat expansions and viral genetic material (or any other organism). Its results are annotated using a variety of databases including ClinVar [11], Exac [12], dbSNP [13] and dbNSFP [14], made available for a local deployment of the gene.iobio platform for an on-the-fly visualisation and user-friendly quality control (QC) reports are generated. DNAscan also allows the user to restrict the analysis to any subregion of the human genome, including the whole exome or a set of genes or gene panel, speeding up the processing time and generating region specific reports. DNAscan is available on GitHub [15] and Docker [16] and Singularity [17] images are also available for fast and reliable deployment.

## Motivation

The pipeline is being developed to meet the needs of the interdisciplinary biomedical research community of the King’s College London Clinical Neuroscience and Health Informatics laboratories. Their current genomics initiatives include a variety of projects ranging from large scale international sequencing projects such as Project MinE and ADNI [18], collecting thousands of whole-genome-sequencing samples, to medical studies designing gene panels for diagnostic purposes. In this context, research and medical workers, with different scientific backgrounds and levels of bioinformatics skills, deal with NGS data constantly. The pipeline requirements were the following:

1. Speed: Some studies for which the pipeline is being used, require the analysis to be concluded within working hours.
2. Usable on personal laptops and desktops: Even if most academic institutions have HPC facilities available for their staff, this does not necessarily mean everything can be processed on them. In many circumstances, many factors, e.g. the informatics skills of the research workers, privacy and ownership policies, technical obstacles, etc., might make using local machines necessary. This is common in a medical research environment.
3. Annotation and visualisation: For non specialised users, e.g. physicians, wet lab biologists, etc., an automatic annotation of the results and user friendly interface for their visualisation is fundamental.
4. Screen for microbial presence: NGS data are widely used to investigate the role of microbes, such as viruses and bacteria, in many diseases. Viral and bacterial metagenomics studies are also common.
5. Specific known repeats: Many repeat expansions have a crucial role in the development of several diseases. E.g. C9orf72 repeat [19] in Amyotrophic Lateral Sclerosis (ALS).
6. Region restricted analysis: Use of shared datasets for heterogeneous research purposes often focuses on subregions of the genome. e.g. screening NGS samples for particular variants or over a disease specific panel of genes.
7. Reproducibility and easy and fast deployment: The pipeline must be easily deployable and results reproducible on any machine. This is to favour both reproducibility of the initial research and collaborations through analysis pipeline sharing.
8. Easy to use: It must be suitable for a wide range of users with different level of informatics expertise.

## RESULTS

### Pipeline description

DNAscan pipeline consists of four stages (Figure 1), Alignment, Analysis, Annotation and Report generation, and can be run in three modes, Fast, Normal and Intensive, according to user requirements. Fast mode minimises the RAM and time required while Normal and Intensive modes improve the variant calling performance by performing an alignment and small indel calling refinement stage respectively.

**Figure 1.**
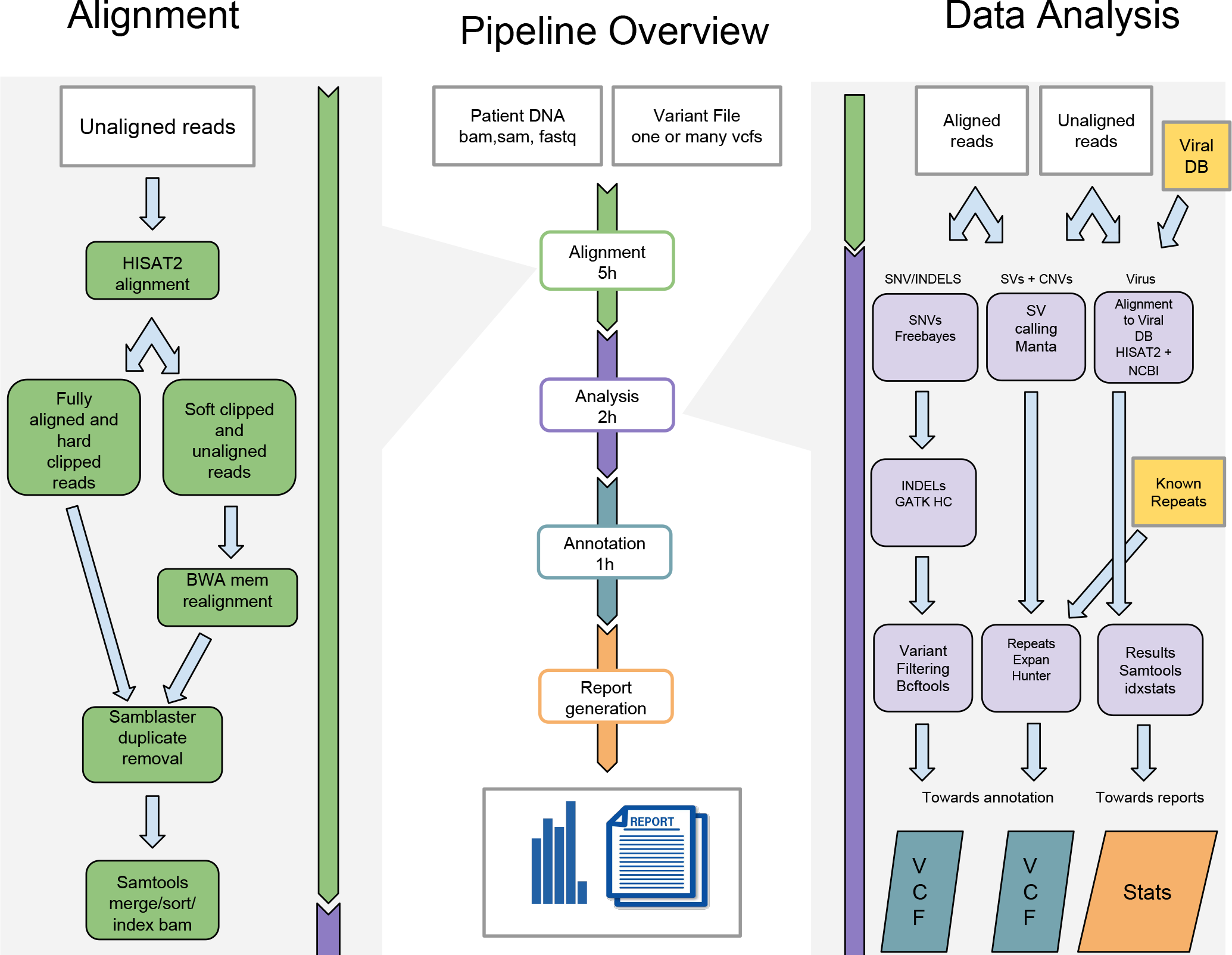
Central panel: Pipeline overview. DNAscan accepts sequencing data, and optionally variant files. The pipeline firstly performs an alignment step (details in the left panel), followed by a customisable data analysis protocol (details in the right panel). Finally, results are annotated and a user friendly QC report is generated. Right panel: detailed description of the post alignment analysis pipeline (Intensive mode). Aligned reads are used by the variant calling pipeline (Freebayes + GATK HC + Bcftools); both aligned and unaligned reads are used by Manta and ExpensionHunter (for which no repeat description files have to be provided) to look for structural variants. The unaligned reads are mapped to a database of known viral genomes (NCBI database) to screen for their DNA in the input sequencing data. Left panel: Alignment stage description. Raw reads are aligned with HISAT2. Resulting soft-clipped reads and unaligned reads are realigned with BWA mem and then merged with the others using Samtools.

The user can tailor the DNAscan pipeline to their needs by performing any subset of the available analyses or restricting them to a subregion of the human genome. e.g. if variants have already been called (this is commonly the case when the sequencing is provided by companies such as Illumina, Novagen, etc) and we are only interested in a particular set of genes, the snv/indel calling steps would be skipped and DNAscan can be used to call SVs only and results visualisation on that set of genes. DNAscan accepts genome regions both as a bed file or a list of gene names. Optionally, the analysis can be restricted to the whole exome.

### Alignment

DNAscan accepts sequencing data both in raw fastq (and its .gz compressed version) and as a Sequence Alignment Map (SAM) file (and its compressed version BAM). HISAT2 [20] and BWA are used to map the reads to the reference genome (Figure 1, left panel). This step is skipped if the user provides ready-aligned data in SAM or BAM formats. HISAT2 is a fast and sensitive alignment program for mapping next-generation sequencing reads to a reference genome. Based on an extension of BWT [21] for graphs [22], HISAT2 implements a large set of small graph FM indexes (GFM) [23] that collectively cover the whole genome. These “local” indexes, combined with several alignment strategies, enable rapid and accurate alignment of sequencing reads. This new indexing scheme is called a Hierarchical Graph FM index (HGFM). Thanks to this novel approach HISAT2 can guarantee a high performance, comparable to state-of-the-art tools, in approximately one quarter of the time of BWA and Bowtie2 [24] (see Results).

Variant calling pipelines based on HISAT2 generally perform poorly on indels [25] (see also Results). To address this issue, DNAscan uses BWA to realign soft-clipped and unaligned reads. This alignment refinement step is skipped if DNAscan is run in fast mode.

Samblaster [26] is used to mark duplicates during the alignment step and Sambamba [27] to sort the aligned reads. Both the variant callers, Freebayes [28] and GATK Haplotype Caller (HC) [29] used in the following step, are duplicate-aware, meaning that they automatically ignore reads marked as duplicate. The user can optionally exclude it from the workflow. This might be necessary in some studies, e.g. when an intensive PCR amplification of small regions is required.

### Analysis

Various analyses are performed on the mapped sequencing data (Figure 1, right panel): SNV and small indel calling is performed using Freebayes, whose reliability is well reported [30, 31]. However, taking advantage of the documented better performance of GATK HC in small indel calling (see also our Results section), we decided to add a customised indel calling step to DNAscan, called Intensive mode. This step firstly extracts the genome positions for which an insertion or a deletion is present on at least one read, and secondly calls indels using GATK HC on these selected positions. The reduced number of positions where this occurs allows for a targeted use of GATK HC, limiting the required computational effort and time. Resulting SNV and small indel calls are finally hard filtered with Bcftools [32] (see methods).

Two illumina developed tools, Manta [33] and Expansion Hunter [34] are used for detecting medium and large structural variants (> 50bp) including insertions, deletions, translocations, duplications and known repeat expansions. These tools are optimised for high speed, and can analyse a 40× WGS sample in about 30 minutes using 8 threads, maintaining a very high performance.

DNAscan also has options to scan the sequencing data for microbial genetic material. It performs a computational subtraction of human host sequences to identify sequences of infectious agents including viruses, bacteria or fungi, by aligning the non-human or unaligned reads to the whole NCBI database [35, 36] of known viral, bacterial or any custom set of microbial genomes and reporting the number of reads aligned to each non-human genome, its length and the number of bases covered by at least one read.

### Annotation

Variant calls are then annotated using Annovar [37]. The annotation includes the use of databases such as ClinVar [11], Exac [12], dbSNP [13] and dbNSFP [38] (more information about how to customise the annotation, e.g. by selecting alternative databases and/or focusing on specific genome regions, are available on GitHub).

### Report generation

Finally, a user friendly, readable and customisable report is generated. Bcftools, Samtools and FastQC (http://www.bioinformatics.babraham.ac.uk/projects/fastqc/) are used to generate statistics on the variants called by the pipeline, the quality of the sequencing data and its alignment. MultiQC [39] is used to wrap up these stats and make them available as html report (see Reports and visualisation utilities). A local deployment of a set of iobio services including the gene.iobio platform (gene.iobio.io) are available for an on-the-fly visualisation of the result variant files (gene.iobio and vcf.iobio) and sequencing data (bam.iobio [40]).

### Software availability

DNAscan is available on GitHub (https://github.com/KHP-Informatics/DNAscan). Docker and Singularity images are also available for fast deployment and reproducibility (see instructions on Github).

### DNAscan benchmarking

Benchmarking every DNAscan component is beyond the aim of this article since a range of literature is available [30, 33, 34, 41]. However, to our knowledge, none exists assessing HISAT2 either for small DNA read mapping or as part of DNA variant calling pipelines. In the following section, we both assess the performance of HISAT2 with BWA and Bowtie2 mapping 1.25 billion WGS reads sequenced with the Illumina Hiseq 2000 and 150 million simulated reads (see Methods), and compare our SNV/indel calling pipeline in Fast, Normal and Intensive modes with the Genome Analysis Toolkit best practices workflow and SpeedSeq [42] over the whole exome sequencing of NA12878. Illumina platinum calls are used as true positives.

We show how DNAscan represents a powerful tool for medical and scientific use by analysing real DNA sequence data from two ALS patients and of HIV infected human cells. For the ALS patients we use both a gene panel of 10 ALS-related genes, whose feasibility for diagnostic medicine has been previously investigated [5], and their equivalent WGS data from the Project MinE sequencing dataset. DNAscan is used to look for SNVs, small indels, SVs, and known repeat expansions. The WGS of an HIV infected human cell sample [43] is used to test DNAscan for virus detection.

### The HISAT2 aligner assessment

To assess the performance of the HISAT2 aligner we used two datasets: 1.25 billion WGS reads of a human WGS DNA sample sequenced with an Illumina Hiseq 2000 (see Methods), and 150 million simulated human reads (see Methods). In this assessment we took into consideration the memory footprint (RAM), the time needed to complete the alignment, percentage of reads mapped to the reference genome, and the percentage of uniquely mapped reads and properly paired reads. The performance of HISAT2 was compared with the BWA aligner (mem algorithm [44]) and Bowtie2, which are two of the most widely used small reads aligners. Table 1 shows the results from this test.

**Table 1:** Alignment assessment results. HISAT2, BWA and Bowtie2 were tested on 150 million simulated Illumina paired end human reads and 1.250 billion real Illumina paired end human reads. For the three aligners on the two dataset the table shows the time taken, their memory fingerprint and the percentage of aligned-one-or-more-times reads, aligned-only-once reads and properly pared. All tests were run using 4 threads.

|  | Simulated reads |  |  | Real reads |  |  |
| --- | --- | --- | --- | --- | --- | --- |
|  | HISAT2 | BWA-MEM | BOWTIE2 | HISAT2 | BWA-MEM | BOWTIE2 |
| Number of reads (Millions) | 150 | 150 | 150 | 1250 | 1250 | 1250 |
| Time (minutes) | 32 | 130 | 115 | 245 | 1167 | 1070 |
| Memory fingerprint (Gigabytes) | 4.2 | 6.6 | 3.8 | 4.2 | 9.1 | 3.8 |
| Aligned reads (%) | 99.82 | 100 | 99.98 | 90.14 | 96.22 | 93.89 |
| Uniquely aligned reads (%) | 97.7 | 100 | 99.98 | 86.17 | 86.80 | 75.14 |
| Properly paired reads (%) | 99.46 | 100 | 52.48 | 85.61 | 95.60 | 62.55 |

On this real dataset, HISAT2 uses 4.2 gigabytes of RAM, slightly larger than Bowtie2 (3.8Gb RAM), while BWA has the biggest memory usage (9.1Gb RAM). In terms of speed, HISAT2 completes the mapping in 4 hours while both the other aligners take about 4-5 times longer (19 hours 27 minutes for BWA and 17 hours 50 minutes for Bowtie2). In terms of percentage of uniquely mapped reads, HISAT2 closely compares with BWA (86.17% and 86.80% respectively) outperforming Bowtie2 (75.14%).

On the simulated dataset, all aligners perform well, however HISAT2 is over 4 times faster than the others, although uniquely aligning slightly fewer reads (97.75% versus BWA-MEM 100%). These results highlight how HISAT2 performs comparably to BWA and Bowtie2 while keeping a low memory footprint (4.2Gb RAM) and the highest speed (over 4 times faster than the other aligners on the real dataset). All tests were run using 4 threads on a machine with 16GB RAM and two Intel Xeon E5-2670 processors.

### Variant calling assessment

To assess the performance of the DNAscan variant calling pipeline with the GATK best practice workflow (GATK BPW) and SpeedSeq, we used the exome of the well studied NA12878 sample and the Illumina platinum calls [45] as a gold standard (our set of true calls). GATK BPW consists of using the BWA aligner, the removal of duplicates with Picard, a base recalibration step and variant calling with GATK-HC. The SpeedSeq pipeline uses BWA for alignment, Sambamba and Samblaster to sort reads and to remove duplicates, and Freebayes for variant calling. Considering the overlap in the software used by DNAseq and SpeedSeq, assessing their performance is therefore of interest. Figure 2A shows the results from this test. DNAscan in Fast mode performs comparably with both the GATK BPW and the SpeedSeq on SNVs. Their F-measure (F_m_), a harmonic mean of precision and sensitivity defined in equation 1 (see Methods), is 0.923, 0.911 and 0.928 respectively.

**Figure 2:**
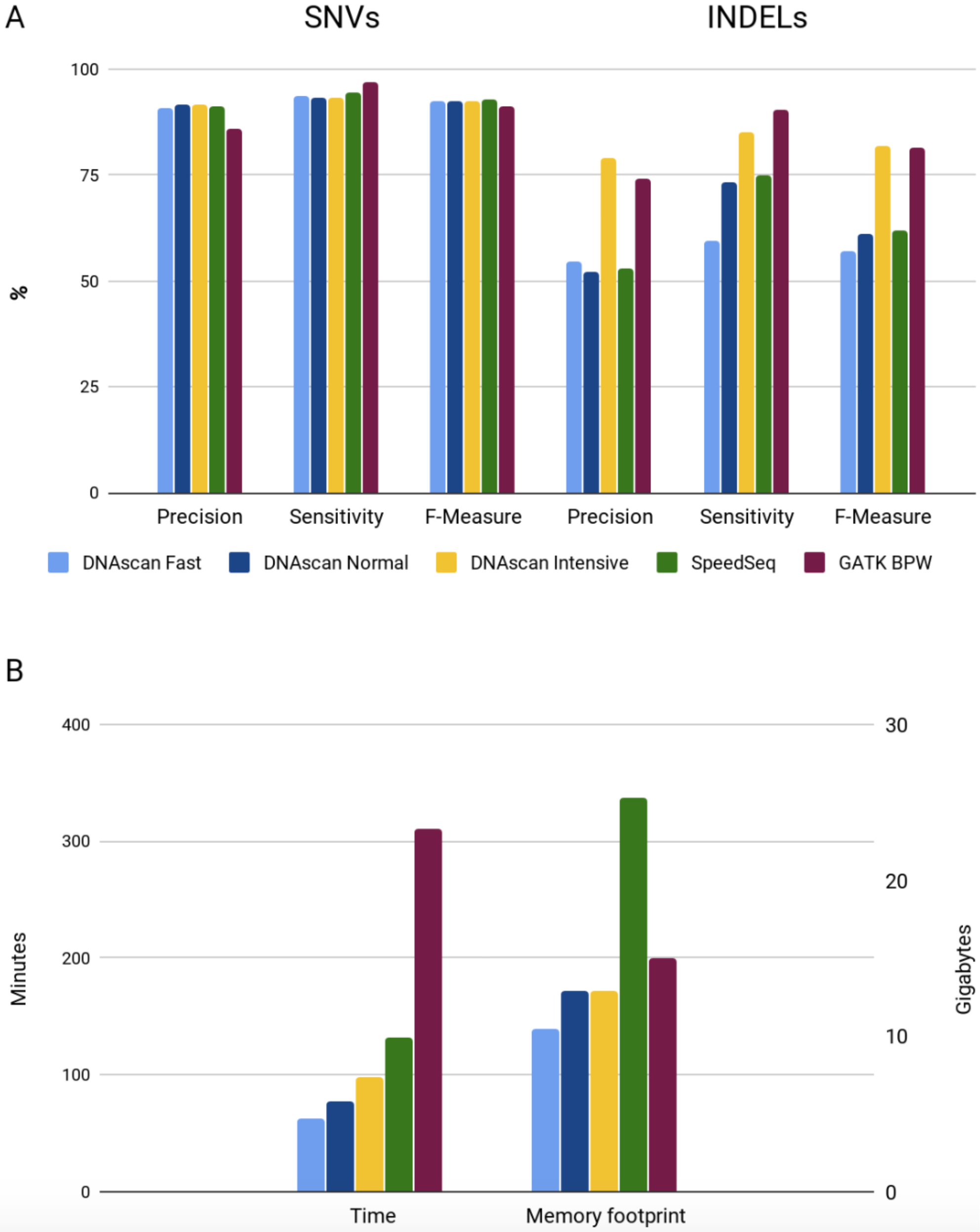
Variant calling assessment. Graph A shows the precision, sensitivity and F-measure of DNAscan in Fast, Normal and Intensive mode, SpeedSeq and GATK best practice workflow (BPW) in calling SNVs and small indels over the whole exome sequencing of NA12878. Illumina platinum calls were used as true positives. The first three columns show the results for SNVs and the last three columns for indels. Graph B shows the time needed and the memory fingerprint for the same pipelines.

For indels, DNAscan in Fast mode performs poorly (F_m_= 0.570). In Normal mode DNAscan shows improvements, reaching an indel calling precision and sensitivity comparable to SpeedSeq (F_m_ equal to 0.610 and 0.620 respectively). The better performance of the Normal mode is driven by a major increase in sensitivity, which reaches 0.734 from 0.596. However, GATK BPW outperforms SpeedSeq on indels (GATK BPW F_m_ = 0.815). DNAscan, in Intensive mode, can perform comparably to GATK BPW also on indels with an F_m_ of 0.820.

Figure 2B shows a comparison of the time needed by the tested pipelines and their memory usage. DNAscan in Fast mode completes the analysis in just 63 minutes while SpeedSeq takes over twice the time (132 minutes) and GATK BPW 5 times longer (310 minutes). DNAscan in both Normal and Intensive mode completes the analysis in a reasonable time (77 and 98 minutes respectively). In terms of memory, DNAscan uses as little as 10GB of RAM in Fast mode, 12GB in Normal and Intensive mode, while GATK BPW 15GB and SpeedSeq over 25GB.

### ALS Miseq and Whole-Genome-Seq test cases

Using DNAscan in Fast mode, we analysed real DNA sequence data from two ALS patients (case A and case B). Case A carries a non-synonymous mutation in the *FUS* gene [46] (variant C1561T, AA change R521C, variant dbSNP id rs121909670 [47]) known to be a cause of ALS (ClinVar id RCV000017609.27,RCV000017611.25). Case B is a confirmed *C9orf72* expansion carrier (see Methods), this repeat expansion is known to be strongly associated with ALS [48]. A panel of 10 ALS related genes was sequenced with the Illumina Miseq platform for both cases, while 40x WGS data was generated with the Illumina Hiseq 2000 for case B only. The Miseq gene panel was designed and tested for diagnostic purposes [3] (see Methods). For these 10 genes (*BSCL2*, *CEP112*, *FUS*, *MATR3*, *OPTN*, *SOD1*, *SPG11*, *TARDBP*, *UBQLN2*, and *VCP*), the full exon set was sequenced, generating over 825,000 222-base-long paired reads. The WGS sample (paired reads, read length = 150, average coverage depth = 40) was sequenced as part of Project MinE sequencing dataset. For both the samples, SNVs, indels, and SVs were called. For the WGS sample, DNAscan also looked for the *C9orf72* repeat.

For the WGS sample we ran DNAscan on the whole genome. However, both for practical reasons and to simulate a specific medical diagnostic interest, we focus our analysis report on the 126 ALS related genes reported on the ALSoD webserver [49] (http://alsod.iop.kcl.ac.uk/misc/dataDownload.aspx#C5).

Frontotemporal Dementia (FTD) is a neurodegenerative disease which causes neuronal loss predominantly involving the frontal or temporal lobes. Considering its genetic and electrophysiological overlap with ALS [50, 51] we report variants linked to FTD as well as to ALS in the following results

Table 2 shows the results from this analysis. For the Case A Miseq DNA gene panel, DNAscan detected 13 SNVs reported to be related to amyotrophic lateral sclerosis and 4 to FTD on ClinVar, 6 non-synonymous variants and 6 variants with a deleteriousness CADD phred score [52] equal to or higher than 13, meaning that they are predicted to be in the top 5% most deleterious substitutions. Finally, the known pathogenic *FUS* SNV rs121909670 was detected. No SVs were found. The whole analysis was performed in ~30 minutes using 4 threads. Since no ALS related repeat expansions are known in these genes, DNAscan was not used to look for any repeat expansions in this analysis.

**Table 2.** Analysis of the ALS patients’ WGS and Miseq data.

|  | Miseq |  | Miseq<br>∩<br>WGS<br>Case B | WGS |  |
| --- | --- | --- | --- | --- | --- |
|  | Case A | Case B |  | gene panel<br>Case B | ALS genes<br>Case B |
| Analysis time (minutes) | 30 | 30 | --- | --- | 460 |
| Data size (MBs) | 40 | 40 | --- | --- | 70,000 |
| N. of ALS-related variants | 13 | 11 | 11 | 11 | 33 |
| N. of FTD-related variants | 4 | 1 | 1 | 1 | 3 |
| N. of non-synonymous variants | 6 | 7 | 3 | 3 | 64 |
| N. of variants with CADD>13 | 6 | 9 | 4 | 4 | 748 |
| N. long insertions | 0 | 0 | 0 | 0 | 1 |
| N. long deletions | 0 | 0 | 0 | 0 | 3 |
| N. Duplications | 0 | 0 | 0 | 0 | 1 |
| N. Inversions | 0 | 0 | 0 | 0 | 0 |
| *C9orf72* expansion | --- | --- | --- | --- | Yes |
| rs121909670 | Yes | --- | --- | --- | --- |

On the WGS data of Case B, for the selected 126 genes, DNAscan identified 33 SNVs reported to be related to amyotrophic lateral sclerosis and 3 to frontotemporal dementia on ClinVar, 64 non-synonymous variants, 748 variants with a deleteriousness CADD phred score equal to or higher than 13, one 60-base-pair long insertion, 3 over 100,000-base-pair long deletions and 1 tandem duplication. DNAscan was also able to detect the known *C9orf72* expansion.

Table 2 also shows the WGS findings restricted to the same regions sequenced with Miseq (table 2 - WGS panel genes column). Considering these results together with the intersection between the WGS and Miseq results of the same ALS patient, thus case B (Miseq ∩ WGS), we can see how all the variants reported to be linked to ALS/FTD on ClinVar were also detected in the WGS data and no novel variants, among the classes considered in table 2, were spotted. The whole analysis was performed in 8 hours using 8 threads.

### Virus scanning

We used DNAscan to detect the presence of viral genetic material in a whole genome sequencing sample of HIV infected human cells (see Methods). The DNA sequencing data was produced using the Illumina Hiseq 2000 sequencer generating about 350 million 95-base-long paired reads (see Methods). Following the well-established approach of computational subtraction of human host sequences to identify sequences of infectious agents like viruses [53], the human reads (91%, Figure 4a) were subtracted by mapping the sequencing data to the reference human genome using HISAT2. To screen our sequencing sample for the presence of known viral DNA, HISAT2 was then used to map the unmapped reads from the initial mapping phase of the pipeline (9%, Figure 4a) to all the viral genomes available on the NCBI virus database.

**Figure 4.**
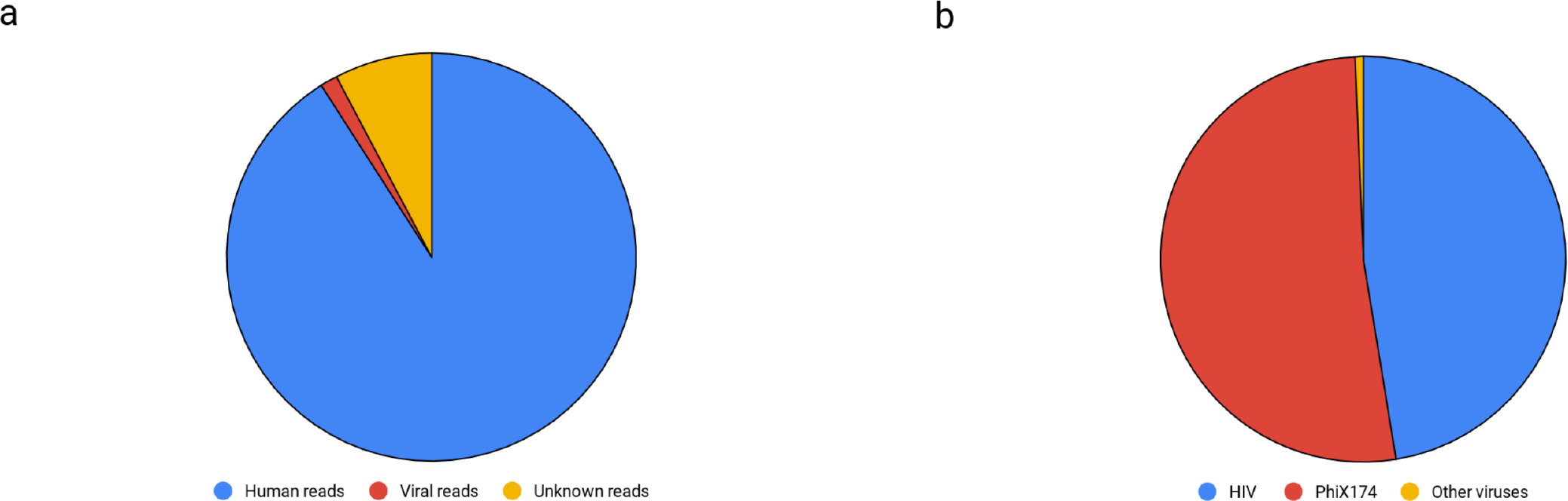
Pie chart a shows the proportion of human reads (blue), viral reads (red) and unknown reads (yellow). Pie chart b shows the proportion for viral reads belonging to HIV (blue), PhiX174 (red) and to other viruses (yellow). Human reads are defined as reads which aligned to the human reference genome, viral reads as the reads which did not align to the human reference genome but aligned to at least one of the NCBI viral genomes and unknown reads as the reads which did not align neither to the human nor to any viral reference genomes.

Figure 4a shows in descending order, the 20 viral genomes to which the highest number of reads were mapped. They show both the presence of HIV DNA and bacterial DNA in our sample. Indeed, 4,412,255 reads mapped to the human immunodeficiency virus (NCBI id NC_001802.1) and only the Escherichia virus phiX174 (NCBI id NC_001422.1), a bacterial virus, presented a comparable number of reads (4,834,017 reads). This phage is commonly found in Illumina sequencing protocols [54] probably because of transfer from gut microbes into blood.

Figure 5b shows a logarithmic representation of the number of reads aligned to the viral genomes, highlighting a smaller (3-4 orders of magnitude) number of reads belonging to other viruses. The disproportion between the presence of the first two hits (phiX174 and HIV) and the rest of the viruses is also shown in Figure 4b. Discussing this heterogeneous viral genetic material in this sample is beyond the aim of this article. The complete results with the list of the whole set of viruses (120 viruses) for which at least one read was aligned can be found on Github (https://alfredokcl.github.io/sample_report/Virus_test.pdf). The whole screening was performed by DNAscan using 4 threads in ~2 hours.

**Figure 5.**
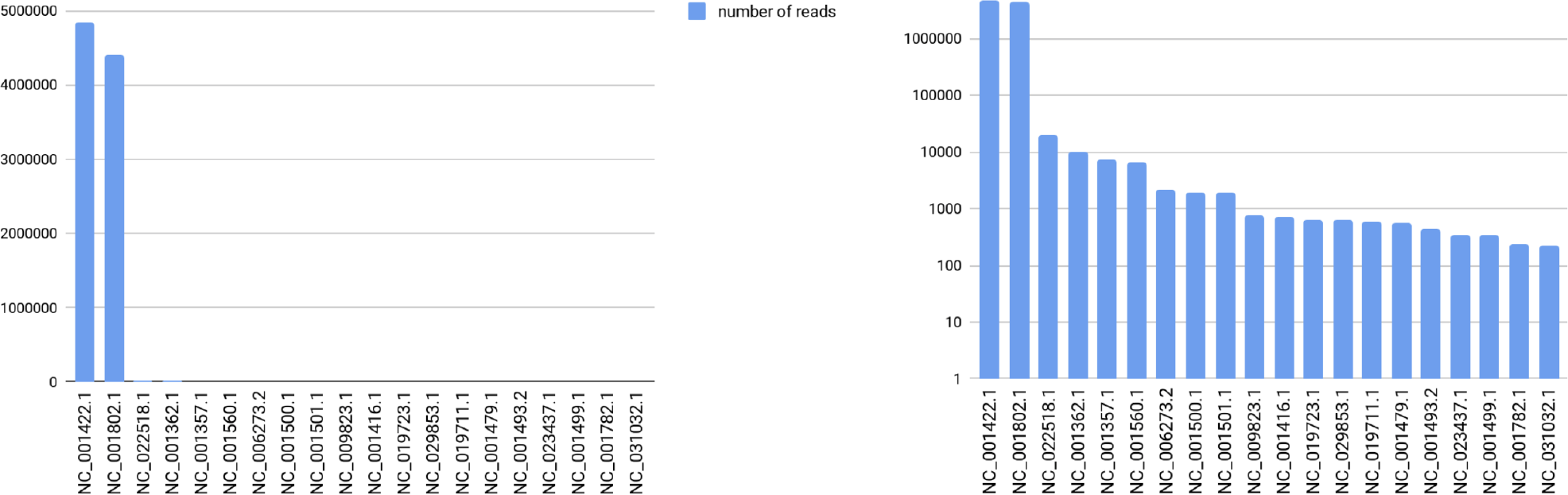
The reads which HISAT2 failed to align to the human reference genome were aligned to the whole NCBI database of viral genomes. In the graphs we plotted numbers of aligned reads in linear (a) and logarithmic (b) scale, for the 20 viral genomes to which the highest number of reads was aligned.

### Reports and visualisation utilities

DNAscan produces a wide set of QC and result reports and provides utilities for visualisation and interpretation of the results.

MultiQC is used to wrap up and visualise QC results of the sequencing data (FastQC), its alignment (Samtools) and variant calls (Bcftools). An example is available at the following link: https://alfredokcl.github.io/sample_report/multiqc_report.html

A tab delimited file including all variants found within a selected region is also generated. This report would include all annotations performed by Annovar in a format that is easy to handle with any Excel-like software by users of all levels of expertise. An example is available at the following link: https://alfredokcl.github.io/sample_report/sample_variant_report.txt

Three iobio services are locally provided with the pipeline allowing for the visualisation of the alignment file (bam.iobio [40]), the called variants (vcf.iobio) and for a gene based visualisation and interpretation of the results (gene.iobio)

These utilities are also available for evaluation at the following links:

http://bam.iobio.io
http://vcf.iobio.io
http://gene.iobio.io

## Discussion

We have presented DNAscan, an extremely fast and computationally efficient pipeline for the analysis, annotation and visualisation of NGS DNA sequencing data. DNAscan can analyse 40× WGS data in 8 hours and WES data in one hour on a mid-range commercial computer. DNAscan also provides utilities for user friendly visualisation and interpretation of its output.

After the assessment showing a positive outcome of HISAT2 versus BWA and Bowtie2, we showed the performance of DNAscan is comparable to the widely used GATK BPW. Its three running modes (Fast, Normal and Intensive) allow the user to tailor the pipeline to their needs, while reducing the time and RAM needed compared to the current GATK BPW and SpeedSeq.

We also reported a few use cases such as the analysis of Miseq and WGS data of an ALS case for diagnostic purposes and the virus screening of HIV infected human cells. In the ALS test we showed how with both technologies DNAscan detected a range of reported ALS related variants in a reasonable time (half an hour for the Miseq panel and 8 hours for the WGS) reporting the presence of both the *C9orf72* expansion and rs121909670. In the HIV test, DNAscan detected the expected viral presence by finding both the HIV virus and a phage commonly present in Illumina NGS DNA data.

Other DNA NGS data analysis pipelines exist. Omictools currently lists 101 such tools. Most of which do not cover the whole data analysis, annotation and visualisation process and are computationally more intensive. Among these SpeedSeq and GATK BPW are two of the most popular. SpeedSeq is a framework for fast genome analysis and interpretation. Analysing a 40× WGS sample with SpeedSeq would take 10 hours on a machine with 32 CPUs and 126Gb of RAM. GATK BPW is a pipeline designed and developed by the Broad institute (https://www.broadinstitute.org) and aims to provide the community with best practice standard pipelines and software for the analysis of NGS data. At present it can be used for SNV and small indel calling. Analysing a WGS sample with the GATK BPW would take about 24 hours on a machine with 32 CPUs and 126Gb of RAM. Compared to these pipelines, DNAscan requires a substantially lower computational effort, provides user friendly utilities for the visualisation and interpretation of results and allows the user to screen the NGS data for microbial genetic material and detect known repeat expansions. Taking into consideration the well reported involvement of microbes and the role of repeat expansions in several diseases of genetic interest [55, 56], both these analyses are valuable research tools.

Cloud computing and storage services offer practically unlimited computational power and storage. However, this has a cost, and its optimisation, in particular for large scale sequencing projects, is of primary importance. Amazon Web Services (AWS) is one of the most popular cloud computing services. Performing the alignment, variant calling and annotation using DNAscan Fast mode on an EC2 instance (https://aws.amazon.com/ec2/pricing/on-demand/) would cost about $2.96 (8 hours of usage of a t2.2xlarge machine with 8 CPUs). The same analysis using SpeedSeq would cost about $22.28 (10 hours of usage of a c3.8xlarge machine with 32 CPUs). These prices do not take into account the storage, were updated on 28th of January 2017 and take into consideration the cheapest machines available in the US East (Ohio) region matching the pipeline computational requirements proposed by the authors (8 CPUs and 16Gb RAM for DNAscan and 32 CPUs and 128Gb RAM for SpeedSeq [42]).

DNAscan is also available as a Docker and a Singularity image. These allow the user to quickly and reliably deploy the pipeline on any machine where either of these programmes is available. Singularity also allows for the deployment of the pipeline on environments for which the user does not have root permission. This could be particularly useful for users working on shared HPC facilities.

DNAscan is a novel and powerful pipeline which can be used for genetic research as well as medical research. It is suitable for both small and large scale analysis. Covering the whole end-to-end analysis process, from sequencing data in fastq format to results visualisation, generating user friendly reports and providing result navigation utilities. DNAscan is suitable for a wide audience of users, ranging from research and medical workers with basic command line usage knowledge to expert bioinformaticians.

## Methods

### Hardware

The tests using four threads were performed on a single machine with 16GB RAM and an Intel i7-670 processor. The tests using more than four threads were performed on a Dell PowerEdge R630 server.

### Performance profiling

Memory usage was recorded using the following command lines:
*$ nohup bash -c ’while true; do (echo "%CPU %MEM ARGS $(date)" && ps -e -o pcpu,pmem,args – sort=pcpu | cut -d" " -f1-5 | tail) >> ps.log; sleep 5; done’ &*

Peak memory usage was parsed by:
*$ cat ps.log | grep -e $proc_name -e ARGS | awk ’BEGIN{max=0}{ if (SUM>max) max=SUM; if ($3=="ARGS") print SUM, SUM=0; else SUM+=$2}END{print max}'*

### Variant calling assessment

To assess the performance of DANscan in calling SNVs and indels, we used the Illumina Genome Analyzer II whole exome sequencing of NA12878 (ftp://ftp-trace.ncbi.nih.gov/1000genomes/ftp/technical/working/20101201_cg_NA12878/NA12878.ga2.exome.maq.raw.bam). Illumina platinum calls (ftp://platgene_ro@ussd-ftp.illumina.com/) were used as true positives.

GATK BPW calls were generated using default parameters and following the indications on https://software.broadinstitute.org/gatk/ (https://software.broadinstitute.org/gatk/best-practices/bp_3step.php?case=GermShortWGS) for germline snvs and indels calling. These include the Pre-processing (https://software.broadinstitute.org/gatk/best-practices/bp_3step.php?case=GermShortWGS&p=1) and variant discovery steps for single sample, i.e. skipping the Merge and Join Genotype steps (https://software.broadinstitute.org/gatk/best-practices/bp_3step.php?case=GermShortWGS&p=2).

SpeedSeq calls were generated running the “align” and “var” commands as described on Github (https://github.com/hall-lab/speedseq)

RTG Tools [57] (“vcfeval” command) was used to evaluate the calls (https://github.com/RealTimeGenomics/rtg-tools).

F-measure, Precision and Sensitivity were defined as in Equation 1.

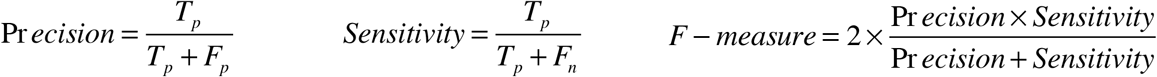

Equation 1: Precision, Sensitivity and F-measure definitions. T_p_ is true positives, F_p_ false positives and F_n_ false negatives.

### *C9orf72* repeat primed PCR

*C9orf72* gene Hexanucleotide Expansions were determined using a Repeat Primed PCR (RP-PCR), previously published by DeJesus-Hernandez et al. [58].

### Simulated reads

150,000,000 Illumina 100-bases-long paired-end human reads were generated using pIRS [59] with default parameters and hg19.

### Whole-Genome-Sequencing data of ALS patients

Venous blood was drawn from patients and controls from which genomic DNA was isolated using standard methods. We set the DNA concentrations at 100ng/ul as measured by a fluorometer with the PicoGreen^®^ dsDNA quantitation assay. DNA integrity was assessed using gel electrophoresis. All samples were sequenced using Illumina’s FastTrack services (San Diego, CA, USA) on the Illumina Hiseq 2000 platform. Sequencing was 150bp paired-end performed using PCR-free library preparation, and yielded ~40× coverage across each sample.

### Miseq ALS gene panel

The ALS gene panel was designed using Illumina TruSeq Custom Amplicon and implemented on an Illumina Miseq platform. This utilises polymerase chain reaction amplicon-based target enrichment and screens for variants in 10 ALS disease genes: *BSCL2*, *CEP112*, *FUS*, *MATR3*, *OPTN*, *SOD1*, *SPG11*, *TARDBP*, *UBQLN2*, and *VCP*. For these genes full exon sequencing was examined.

### Whole-Genome-Sequencing of HIV infected human cells

Genomic libraries were prepared using the TruSeq^®^ DNA Sample Prep kit V2 (Illumina) following the manufacturer’s instructions. Briefly, 1 μg of genomic DNA was sheared with the Covaris 2 system (Covaris). The DNA fragments were then end-repaired, extended with an ‘A’ base on the 3′ end, ligated with indexed paired-end adaptors and PCR amplified. PCR amplification was carried out as follows: initial denaturation at 98°C for 30 sec, followed by 8 cycles consisting of 98°C for 10 sec, 60°C for 30 sec and 72°C for 30 sec, then a final elongation at 72°C for 5 min. Four different genomic libraries were pooled and sequenced in one lane of an Illumina Hiseq 2000 sequencer using a 2 × 95bp paired end indexing protocol. Demultiplexed fastq files were obtained for each sample using the Illumina CASAVAv1.8.1 software. The complete high throughput sequencing dataset was downloaded from the Sequence Read Archive (SRA) [60] under accession number SRA056122.

### Viral database

In this paper DNAscan makes use of the whole non-redundant NCBI database of complete viral genomes (9334 genomes). These can be downloaded both as a multi sequence fast file, together with its fai index and HISAT2 index from our GitHub repository (link) and directly from the NCBI database ftp server (ftp.ncbi.nlm.nih.gov/refseq/release/viral)

DNAscan can also be used to screen for the DNA of other organisms including bacteria or fungi in which case the user can download the preferred database from the NCBI ftp server or from our GitHub where the corresponding index files can also be found (e.g. ftp.ncbi.nlm.nih.gov/refseq/release/bacteria or ftp.ncbi.nlm.nih.gov/refseq/release/)

### List of tools used in this work

- Genome Analysis Toolkit 3.8
- BWA 0.7.15
- Picard 2.2.1
- Samtools 1.5
- HISAT2 2.1.0
- Bcftools 1.5
- RTG Tools 3.6.2
- Python 3.5
- MultiQC 1.2
- FastQC v0.11.7
- Docker 1.7.1
- Docker-compose 1.4.2
- Freebayes 1.0.2
- ExpansionHunter 2.0.9
- Manta 1.0.3
- Annovar "Version: $Date: 2016-02-01 00:11:18 −0800 (Mon, 1 Feb 2016)”
- Bedtools2 2.25

